# Clustering of HR+/HER2- breast cancer in an Asian cohort is driven by immune phenotypes

**DOI:** 10.1101/2023.12.07.570545

**Authors:** Jia-Wern Pan, Mohana Ragu, Wei-Qin Chan, Siti Norhidayu Hasan, Tania Islam, Li-Ying Teoh, Suniza Jamaris, Mee-Hoong See, Cheng-Har Yip, Pathmanathan Rajadurai, Lai-Meng Looi, Nur Aishah Mohd Taib, Oscar M. Rueda, Carlos Caldas, Suet-Feung Chin, Joanna Lim, Soo Hwang Teo

## Abstract

Breast cancer exhibits significant heterogeneity, manifesting in various subtypes that are critical in guiding treatment decisions. This study aimed to investigate the existence of distinct subtypes of breast cancer within the Asian population, by analysing the transcriptomic profiles of 934 breast cancer patients from a Malaysian cohort. Our findings reveal that the HR+/HER2-breast cancer samples display a distinct clustering pattern based on immune phenotypes, rather than conforming to the conventional luminal A-luminal B paradigm previously reported in breast cancers from women of European descent. This suggests that the activation of the immune system may play a more important role in Asian HR+/HER2-breast cancer than has been previously recognized. Analysis of somatic mutations by whole exome sequencing showed that counter-intuitively, the cluster of HR+/HER2-samples exhibiting higher immune scores was associated with lower tumour mutational burden, lower homologous recombination deficiency scores, and fewer copy number aberrations, implicating the involvement of non-canonical tumour immune pathways. Further investigations are warranted to determine the underlying mechanisms of these pathways, with the potential to develop innovative immunotherapeutic approaches tailored to this specific patient population.

## Introduction

Breast cancer is a highly heterogeneous disease, characterized by diverse subtypes that have important implications in guiding treatment^1–4^. These distinct subtypes encompass variations in clinical presentation, molecular profiles, and response to treatment, highlighting the complex nature of breast cancer. The classification of breast cancer subtypes has provided valuable insights into disease prognosis and treatment selection, enabling more personalized and targeted therapeutic approaches.

Currently, there are two generally accepted methodologies to classify breast cancer subtypes: PAM50^1,2,5^ and Integrative Clusters^3,4,6^. Under the PAM50 classification scheme, breast cancer is classified into four main subtypes according to gene expression-based clusters: luminal A, luminal B, Her2-enriched, and basal-like, as well as an additional small group usually labelled as “normal-like”. These categories roughly correspond to the biomarker- and treatment-based clinical subtypes of HR+/HER2-, HR+/HER2+, HR-/HER2+, and triple negative breast cancer (TNBC). The Integrative Cluster classification scheme, on the other hand, integrates both transcriptomic and copy number data to classify breast cancer into 11 subtypes with different driver mutations. These subtypes include the HER2+-associated IntClust 5, the TNBC-associated IntClust 10, as well as subtypes with low copy number variation (IntClust 4+ and 4-).

However, it is important to recognize that the existing subtypes were predominantly derived from studies conducted in populations of European descent. Given the clinical utility of molecular subtyping, there is a growing need to investigate breast cancer subtypes in different ethnic populations to account for potential variations in genetic, environmental, and lifestyle factors. For example, the various breast cancer subtypes are known to have different associations with reproductive risk factors^7,8^ and BMI^7^, and these lifestyle risk factors are likely to differ substantially between different populations. Additionally, different breast cancer subtypes are associated with different breast cancer risk genetic loci^9–11^, which are also distributed differently in different populations^12^. Importantly, previous studies have found differences in the prevalence of certain subtypes in non-European populations, with the African population having higher numbers of TNBC-associated samples^13^, and the Asian population having a higher prevalence of Her2-enriched^14^ or luminal B^15^ samples.

The heterogeneity in clinicopathological features across different populations suggests the possibility that intrinsic molecular subtypes differ across different populations as well. For example, breast cancers from the Nigerian population have a high prevalence of homologous recombination deficiency (HRD), *TP53* mutations, and structural variation indicative of a more aggressive biology compared to tumours from Western populations^13^. It has also been identified previously that Asian breast cancers exhibit higher immune scores compared to the Western population^14,15^. These findings suggest potential differences in the underlying biology of breast cancer in different ethnic groups and highlight the importance of population-specific studies across diverse populations.

Investigating breast cancer subtypes in Asians holds significant promise for improving our understanding of the disease and optimizing treatment strategies for this specific population. Additionally, it may provide insights into the underlying genetic and molecular mechanisms that contribute to breast cancer pathogenesis, allowing for the development of more tailored and effective therapeutic interventions. Existing studies which investigated the molecular subtypes underlying breast cancer in Asian populations have reported that molecular subtypes are largely conserved between Asian and Western populations^15–17^. However, the cohort sizes in these studies are relatively small, and some are only focused on a subpopulation, such as young breast cancer, and may not be representative of the entire Asian population. In this study, we explored the possibility of unique subtypes of breast cancer within the Asian population by examining the transcriptomic profiles of breast cancer patients from a relatively large cohort. Our analyses found the HR+/HER2-breast cancer samples in our cohort display a distinct clustering pattern based on immune phenotypes instead of the luminal A-luminal B paradigm. This suggests that the activation of the immune system may play a more important role in Asian HR+/HER2-breast cancer than has been previously recognized, which may have important clinical implications.

## Materials & Methods

### Study cohort

Our study cohort consists of 934 Malaysian women with breast cancer. Patients were recruited to the MyBrCa Genetics study^18^ from the Subang Jaya Medical Centre and the University Malaya Medical Centre between 2012 and 2018. Peripheral blood samples and breast tumour tissues were acquired from each patient. For breast tumour tissues, representative fresh tumour tissues were obtained and frozen during surgical resection of the tumour. These tumour samples were then stored in liquid nitrogen. Tumour samples were then sectioned for DNA and RNA extraction. The top and bottom sections were stained with haematoxylin and eosin and reviewed for tumour content. Tumour samples with an average tumour content of <30% (*n*□=□50) and/or insufficient DNA (*n*□=□8) were excluded from the study. Patients with bilateral breast cancer were also excluded (*n=*14). Patient enrolment and genetic analyses were approved by the Ethics Committee of Subang Jaya Medical Centre (reference no: 201208.1) and the Medical Ethics Committee of the University Malaya Medical Centre (reference no: 842.9) and written informed consent was provided by each patient.

### Nucleic acid extraction and sequencing of tumour and matched normal specimens

For samples which were part of our original MyBrCa tumour cohort publication, DNA and RNA extraction and sequencing were performed as previously published^14^. This study also included an additional 448 samples. For these samples, DNA extraction from blood samples was performed using the Maxwell 16 Blood DNA Purification Kit with a Maxwell 16 Instrument, following standard protocol. For tumour samples, DNA was extracted using the QIAGEN DNeasy Blood and Tissue Kit following standard protocol and quantified using the Qubit HS DNA Assay kit and Qubit 2.0 fluorometer (Life Technologies Inc).

RNA was extracted from tumour samples using the QIAGEN miRNeasy Mini Kit with a QIAcube, according to standard protocol. Quantification of total RNA was performed using a Nanodrop 2000 Spectrophotometer and RNA integrity was established through an Agilent 2100 Bioanalyzer. For both DNA and RNA sequencing (RNA-seq), samples with a concentration above 20.0 ng/μL were selected. Additionally, an RNA integrity number above 7 was required for samples to be selected for RNA-seq.

DNA libraries were produced from 50 ng of genomic DNA using the Nextera Rapid Capture Exome kit (Illumina, San Diego, USA) as per manufacturer’s instructions. Exome capture was achieved in pools of 3 and subjected to paired end 75 sequencing on a NovaSeq platform (Illumina, San Diego, USA) at 40x depth for blood samples and 80x depth for tumour samples. Prior to exome capture, 4 nM pools of DNA libraries from tumour samples were also selected for single end 50 shallow whole-genome sequencing at 0.1x depth.

RNA libraries were prepared from 550 ng of total RNA from tumour samples using the TruSeq Stranded Total RNA HT kit with Ribo-Zero Gold (Illumina, San Diego, USA) as per manufacturer’s instructions, subjected to paired end 75 sequencing on a NovaSeq platform (Illumina, San Diego, USA) at 40x depth.

### Gene Expression Analysis

RNA-seq reads were aligned to the hs37d5 human genome and the ENSEMBLE GrCh37 release version 87 human transcriptome via the STAR aligner (v.2.5.3a)^19^. Gene-level counts were calculated with featureCounts (v. 1.5.3)^20^. Gene-level count matrices for the cohort were transformed into normalized log2 counts-per-million (logCPM) using the voom function from the limma (v. 3.34.9) R package. The transformed matrices were then subtyped according to PAM50 and SCMgene designations using the Genefu package in R (v. 2.14.0).

### Clustering and Classification Analysis

To identify unique subtypes, unsupervised k-means clustering was performed on the gene-level count matrices for the MyBrCa cohort. We also evaluated the use of different numbers of genes as our feature set, by ranking each gene according to the median absolute deviation of each gene within the cohort, and using either the top 1000 genes with the highest median absolute deviation, the top 5000, or all genes. To ensure robustness, an extensively implemented consensus clustering method^21^, with 1000 iterations and 0.9 subsampling ratio, was used to assess clustering stability. Consensus clustering was implemented by the ConsensusClusterPlus function of the R package ConsensusClusterPlus with k-means clustering algorithm using Pearson correlation distance. Our final clustering model used k=12 with the top 5000 genes.

### Hierarchical clustering analysis

Hierarchical clustering was conducted on the gene-level count matrices for the MyBrCa cohort using the hclust package from R with default parameters.

### Shallow whole genome sequencing alignment and copy number aberration (CNA) assessment

Sequenced reads were mapped to the hg19 reference genome using bwa-mem, sorted using samtools and dedupped using picard (http://broadinstitute.github.io/picard). Mapped reads were analysed using QDNAseq^22^ to obtain 100 kb segmented copy number profiles using standard protocol and default parameters. CNAs were called using CGHcall (v 2.40) as implemented in the QDNASeq R package, which calls each segment as normal, copy number gain, copy number loss, amplification or deletion using a mixture model. ENSEMBL hg19 genes with HUGO names were mapped to the segmented copy number calls by their start positions to determine the copy number status for each gene in each sample.

### Profiling the tumour immune microenvironment

Overall immune cell infiltration in the bulk tumour samples was assessed from RNA-seq TPM gene expression scores using ESTIMATE (v. 1.0.13)^23^, gene set variation analysis (GSVA) (v. 1.26) using combined immune cell gene sets from Bindea et al. (2013)^24^ and immune scores from Thorsson et al. (2018)^25^. For each sample, immune features predictive of checkpoint inhibitor immunotherapy was also scored. This was done using IMPRES scores (only 14 out of 15 of the predictive features were available in our datasets) as well as GSVA using the Expanded IFN-gamma gene set^26,27^. The relative abundance of specific immune cell populations was estimated from RNA-seq TPM scores with the CIBERSORT^28^ web tool, as well as through GSVA with individual immune gene sets from Bindea et al. (2013).

### Mutational signatures

The weights of previously reported breast cancer mutational signatures using COSMIC matrices (Single Base Substitutions (SBS) Signatures 1, 2, 3 and 13) were established using deconstructSigs^29^. The proportion of variants associated with each mutational signature was determined only for samples with at least 15 detected single nucleotide variants (SNVs).

### HRD scores

The following measures of HRD were determined as described previously: (1) loss of heterozygosity (LOH), (2) large-scale state transitions (LST), and (3) telomeric allelic imbalance (TAI)^30,31^. Allele-specific copy number (ASCN) profiles on paired normal-tumour BAM files were classified via Sequenza^32^ and utilised to analyse the individual measure scores and HRD-sum scores via scarHRD R package^33^.

### Differential gene expression and functional enrichment analysis

Gene expression was analysed with the DEseq2 package, an R-based open-source software designed to analyse transcriptomic data for differential expression, as previously described^34^. Gene set enrichment analyses (GSEA) was performed to compare clusters using the Hallmark pathways from the Molecular Signatures Database^35^ as well as KEGG pathways. These analyses were performed with default parameter settings using 1,000 permutations and an FDR cutoff of 0.05. Single-sample gene set enrichment analyses (ssGSEA) and gene set variation analyses (GSVA) were also performed for each individual sample for specific Hallmark and KEGG pathway gene sets, including the Hallmark complement, hypoxia, IL6-JAK-STAT3 signalling, inflammatory response, and interferon gamma response pathways, as well as the KEGG cGAS-STING, T-cell receptor signalling, antigen processing and presentation, and TGF-β signalling pathways, using the GSVA package in R.

### Survival analysis

Survival data of patients were obtained from the Malaysian National Registry record of deaths. Survival length was defined as the period between the date of diagnosis of patients until the date of death for deceased patients, or the date when the Malaysian National Registry was last queried for patients assumed to be still alive. Cox proportional hazard models were built using the coxph function from the survival package and plotted using ggadjustedcurves and ggforest functions from the survminer package in R (v. 4.3.1). For comparison of overall survival between MyBrCa clusters, the cluster with relatively large sample sizes and the best survival (Clusters 2) was selected as the reference group, and the p-value from comparison with Cluster 1, which had the worst survival among the larger clusters, was reported. For comparison between HR+/HER2-clusters, Cluster 7 (Group 1) was used as the reference group as comparison with the Group 2 clusters and the p value of the cluster with best survival was reported.

### Statistical analysis

The Wilcoxon test and the Chi-square test were executed for comparisons of variables between categories. Unpaired t-tests were used to compare continuous variables between two groups. All tests were two-tailed and a significance level of *p*=0.05 was used. Statistical analyses were performed using R v4.0. All box and whiskers plots in the main and supplementary figures were constructed with boxes indicating the 25th percentile, the median and the 75th percentile, whiskers showing the maximum and minimum values within 1.5 times the inter-quartile range, and outliers were not shown.

## Results

### Clinical characteristics of the MyBrCa cohort

The MyBrCa tumour cohort comprises of 934 female breast cancer patients of self-reported Malaysian nationality who were sequentially recruited from two Malaysian hospitals, Subang Jaya Medical Centre and Universiti Malaya Medical Centre. This cohort consists of a mix of Chinese, Malay, or Indian ancestry. The clinical characteristics of these patients are shown in Table 1.

**Table 1.** Clinico-pathological characteristics of the study cohort.

|  | <u>Overall</u> | <u>Group 1</u> | <u>Group 2</u> | <u>Statistical</u><br><u>Significance</u> |
| --- | --- | --- | --- | --- |
| **Subjects (n)** | 934 | 61 | 421 |  |
| **Patient age (mean±SD)** | 53.78±11.65 | 50.56±10.08 | 54.16±11.89 | 0.0247 |
| **Clinical subtypes (n(%))** |  |  |  |  |
| **HR-/HER2+** | 138 (14.78) | 5 (8.20) | 5 (1.19) | 0.0004 |
| **HR+/HER2-** | 432 (46.25) | 39 (63.93) | 312 (74.11) |  |
| **HR+/HER2+** | 126 (13.49) | 10 (16.39) | 60 (14.25) |  |
| **TNBC** | 159 (17.02) | 4 (6.56) | 7 (1.66) |  |
| **N/A** | 79 (8.46) | 3 (4.92) | 37 (8.79) |  |
| **TNM Stage (n(%))** |  |  |  |  |
| **0** | 23 (2.46) | 2 (3.28) | 8 (1.90) | 0.7430 |
| **I** | 146 (15.63) | 8 (13.11) | 67 (15.91) |  |
| **II** | 428 (45.82) | 28 (45.90) | 198 (47.03) |  |
| **III** | 270 (28.91) | 20 (32.79) | 114 (27.08) |  |
| **IV** | 40 (4.28) | 2 (3.28) | 24 (5.70) |  |
| **N/A** | 27 (2.89) | 1 (1.64) | 10 (2.38) |  |
| **Grade (n(%))** |  |  |  |  |
| **1** | 27 (2.87) | 3 (4.92) | 22 (5.23) | 0.3310 |
| **2** | 385 (40.45) | 31 (50.82) | 248 (58.91) |  |
| **3** | 420 (44.46) | 22 (36.07) | 113 (26.84) |  |
| **N/A** | 102 (12.22) | 5 (8.20) | 38 (9.03) |  |
| **Histology (n(%))** |  |  |  |  |
| **Ductal Carcinoma** | 800 (85.65) | 48 (76.19) | 361 (82.23) | <0.0001 |
| **Lobular Carcinoma** | 30 (3.21) | 11 (17.46) | 11 (2.51) |  |
| **Other** | 1 (0.11) | 0 (0.00) | 0 (0.00) |  |
| **N/A** | 103 (11.03) | 2 (3.17) | 49 (11.16) |  |

### Identification of Clusters Associated with Subtypes of MyBrCa patients

After sample processing, sequencing, and data processing, we were able to obtain transcriptomic profiles for 934 samples, which we used to conduct a cluster analysis. Using k-means clustering, we iteratively analysed the clustering of these samples across a range of *k* values and number of features (number of genes included – see Methods). The optimum *k* value and feature set was selected by comparing our clustering results with the PAM50 clustering. Given that Her-2 enriched breast cancer is a well validated, biologically distinct subtype with copy number amplification of the ERBB2 gene, we selected a clustering result which grouped Her2-enriched samples consistently into a single cluster to be used for all subsequent analyses. This clustering result had the *k* value of 12 and a feature set of the top 5000 genes with the highest median absolute deviation within the cohort (Figure 1).

**Figure 1.**
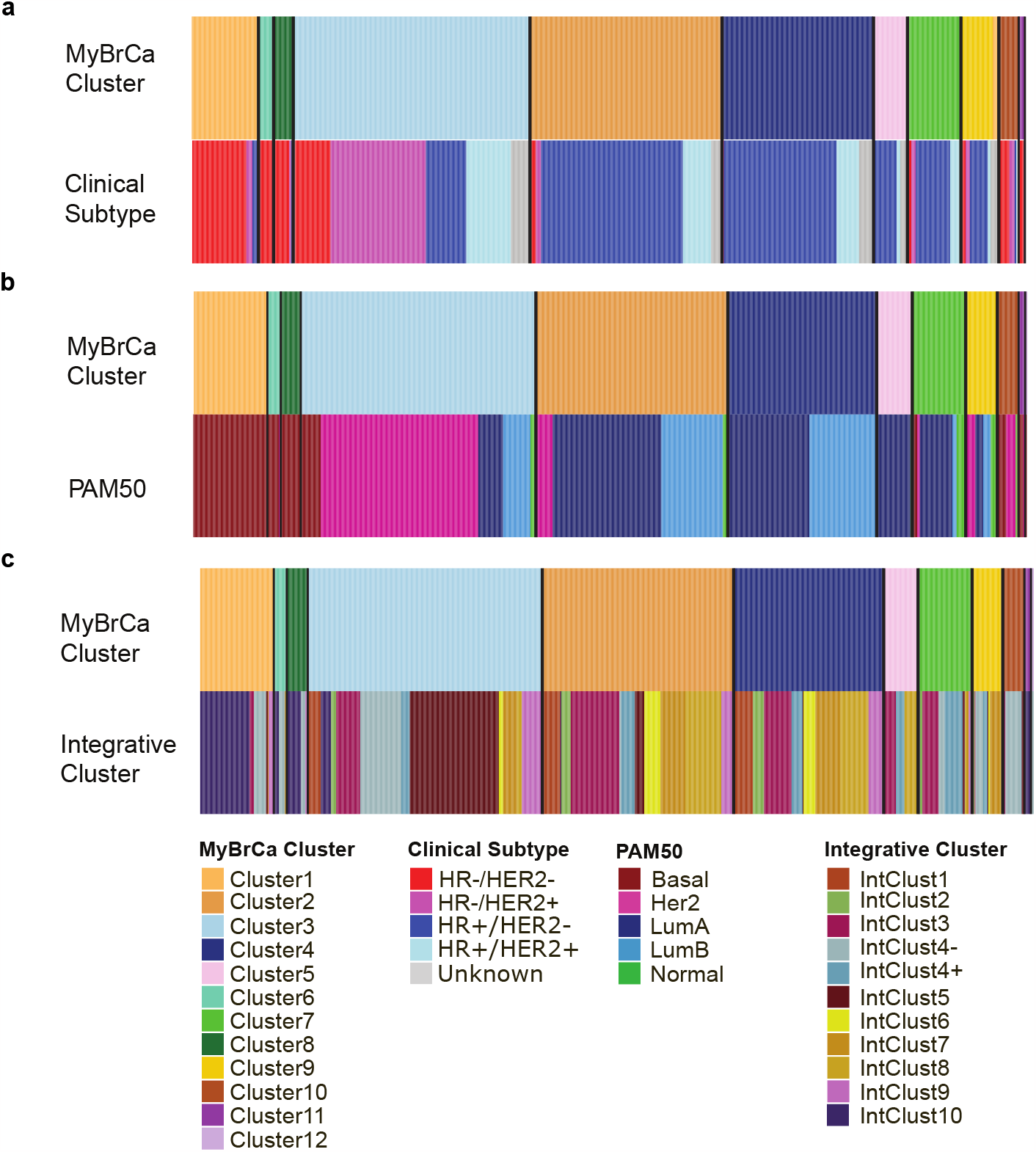
Comparison of MyBrCa clusters with **(a)** clinical subtypes, **(b)** PAM50 and **(c)** Integrative Clusters.

Comparisons with PAM50, IntClust, and clinical subtypes indicated that our clustering results largely support the notion of distinct basal-like/IntClust10/TNBC and Her2-enriched/IntClust5/HER2+ groups of samples (Figure 1). However, our clustering results were very different compared to previous clustering methods for the subtypes of HR+ samples (Figure 1a), suggesting that the grouping of HR+ samples in our cohort may be driven by a different phenotypic paradigm relative to other populations.

### Comparison of pathway expression across MyBrCa Clusters

Next, to investigate the differences between our clusters, we conducted differential gene expression and pathway analyses. We compared each cluster to the other clusters with similar clinical subtypes, with a particular focus on the HR+ clusters. Using GSEA, we found that the HR+ clusters 2, 4, 5, and 7 differed from each other primarily in immune-related pathways such as the complement, inflammatory response, and interferon gamma (IFN-γ) response pathways (Figure 2c, Supplementary Table 1). Pathway enrichment analyses returned similar results, with an over-representation of genes involved in immune-related pathways when comparing between the four HR+ clusters. To follow up on these results, we calculated the scores for several gene expression-based immune-scoring methods, including ESTIMATE, IMPRES, the Ayers expanded IFN-γ gene set, and the immune scores from Thorsson et al. (2018) for each sample (Figure 2b). We found that there was a marked difference for many of these scores between our HR+ clusters, consistent with the notion that the clustering of HR+ samples in our cohort is driven by differences in immune-related phenotypes. We identified a group of clusters with consistently low or intermediate immune scores (Group 2), comprising of Clusters 2, 4 and 5, while Cluster 7 (Group 1) has consistently high immune scores.

**Figure 2.**
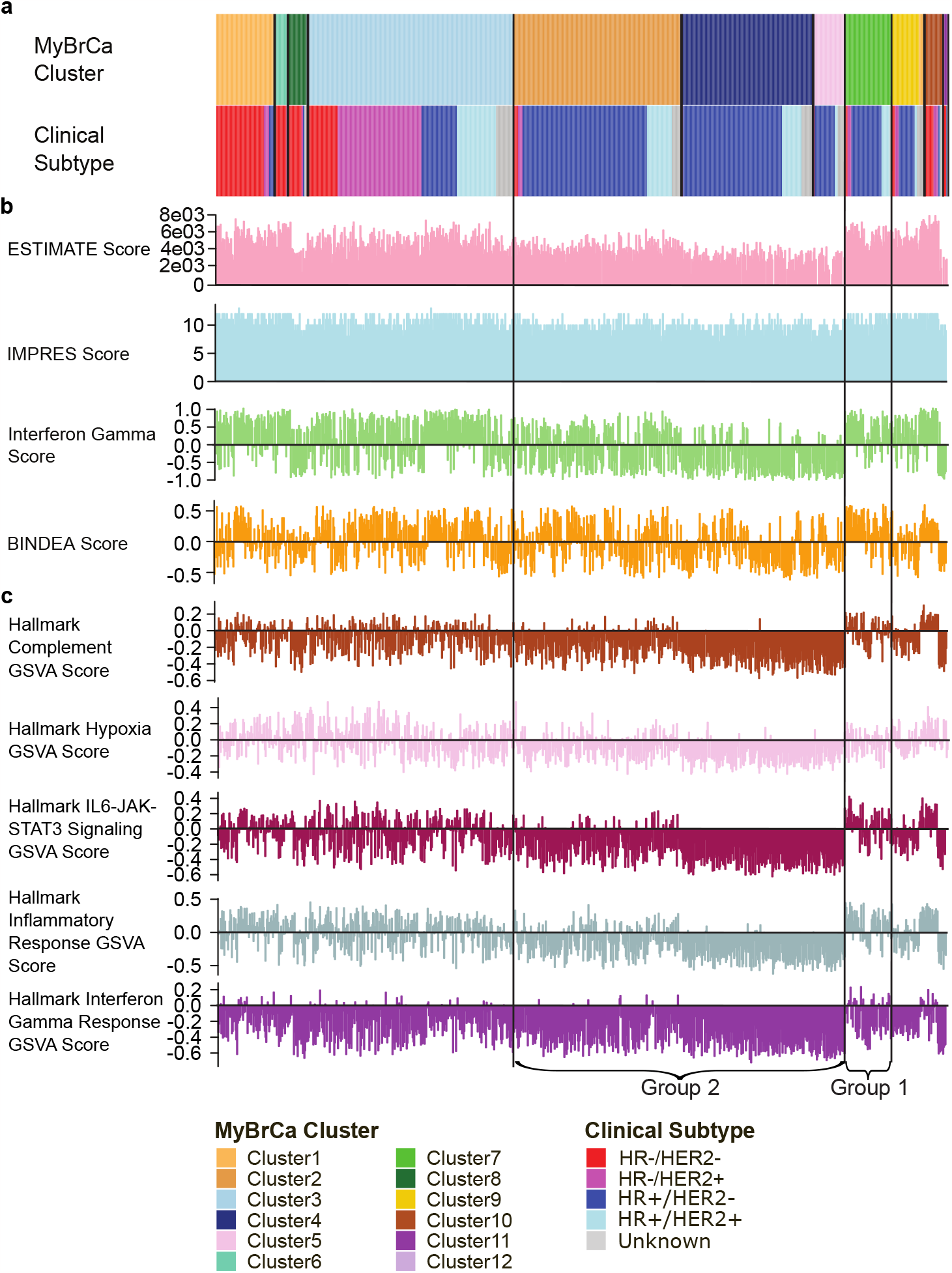
Comparison of MyBrCa Clusters with **(a)** clinical subtypes, **(b)** immune scores and **(c)** Gene Set Variation Analysis (GSVA) scores. HR+ clusters with high immune scores (Group 1) and low immune scores (Group 2) are identified.

We also investigated several KEGG pathways known to mediate IFN-γ, including cGAS-STING, T-cell receptor signalling, antigen processing and presentation, and TGF-β signalling pathways. We compared gene expression for genes in these pathways between our Group 1 and Group 2 samples using GSVA and ssGSEA, and the results indicated that all of these pathways were upregulated in Group 1 as compared to Group 2 (Supplementary Figure 1).

We validated our results by conducting unsupervised hierarchical clustering of transcriptomic profiles of the MyBrCa cohort. Hierarchical clustering of our samples gave similar results to our k-means clustering analyses in that our HR+/HER2-samples were again clustered into two main groups, and the segregation of the two groups was primarily driven by immune-related pathways according to pathway analyses (Supplementary Figure 2, Supplementary Table 2).

### Genomic profiles of HR+/HER2-clusters

Given the growing importance of immunotherapy in cancer treatment, biomarkers to identify a subset of HR+/HER2-breast cancer patients that have an active immune microenvironment may be useful in both clinical and research settings. Since both whole-exome and shallow whole-genome sequencing data are available for the majority of samples in our study cohort, we next compared the mutational and copy number profiles of samples in the high immune scoring HR+/HER2-cluster (Group 1) to those in the intermediate and low immune-scoring HR+/HER2-clusters (Group 2), in order to identify molecular features that may be associated with an active immune microenvironment in a HR+/HER2-breast cancer background. First, we compared the prevalence of known somatic driver mutations in both groups and found that somatic *TP53* mutations were more common in Group 1, but no other driver mutations were significantly different (Figure 3a). Next, we compared the prevalence of mutational signatures and found that the aging-associated mutational signature SBS1 was more prevalent in Group 2 (Figure 3b). Contrary to expectations, samples in Group 1 had on average fewer somatic mutations (Figure 3c) and CNAs (Supplementary Figure 3) compared to Group 2, as well as lower scores for HRD-associated LOH and LST (Figure 3d).

**Figure 3.**
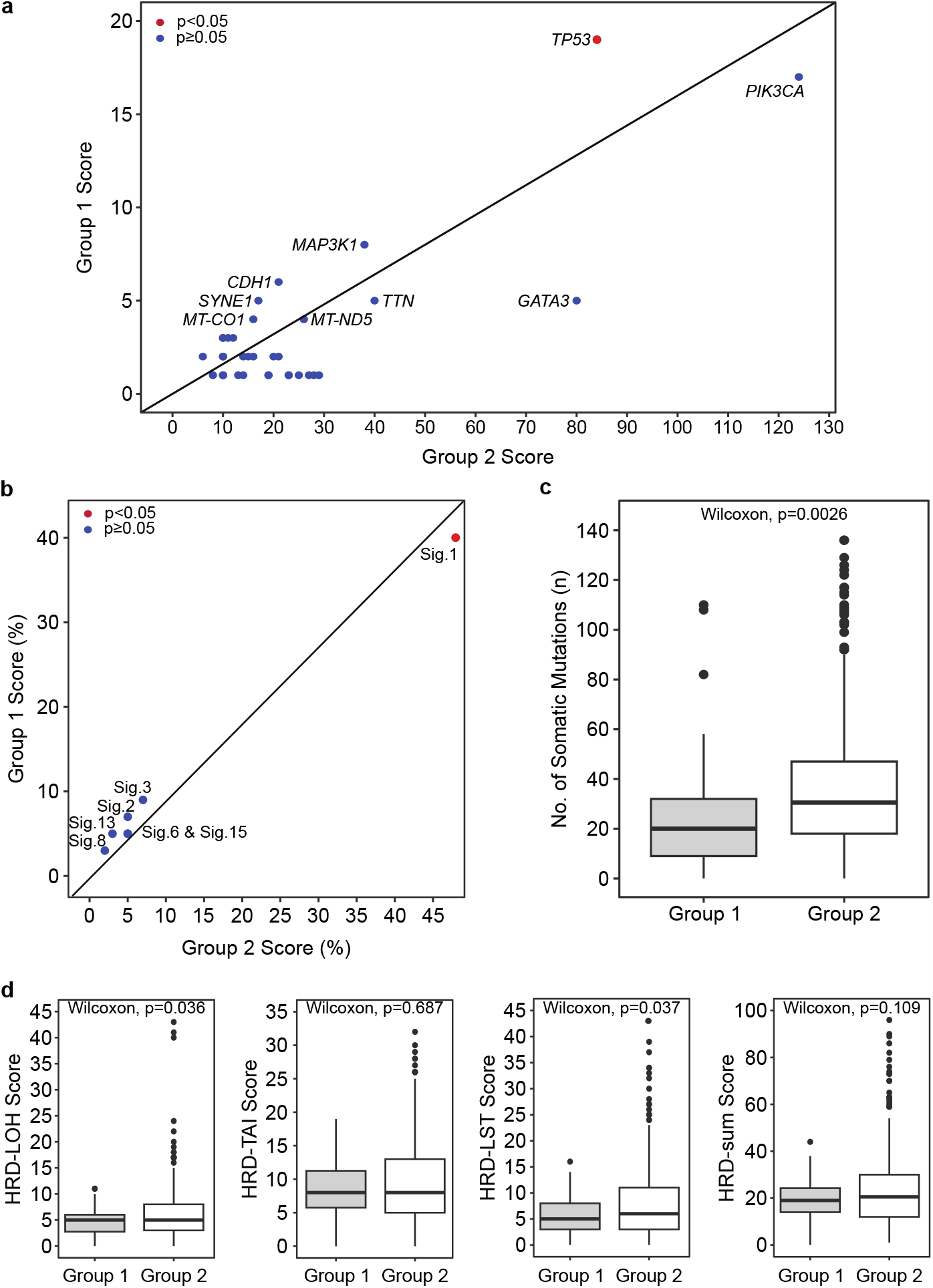
Comparison of characteristics between HR+/HER2-Group 1 and Group 2. **(a)** Somatic driver mutations in Group 1 and Group 2. **(b)** Mutational signatures of Group 1 and Group 2. **(c)** Number of somatic mutations in Group 1 and Group 2. **(d)** Comparison of homologous recombination deficiency (HRD) scores between Group 1 and Group 2. LOH= loss of heterozygosity, LST= large-scale state transitions, TAI= telomeric allelic imbalance. A signifance level of p=0.05 was used.

### Immune profiles of HR+/HER2-clusters

Next, we asked if there was a difference in the composition of immune cells in the tumour microenvironment between the two groups that could be associated with immune activation. We first confirmed that there was a significant difference between the two groups by comparing several general immune score markers and found that Group 1 indeed had significantly higher scores for the combined Bindea gene set, Ayers expanded IFN-gamma gene set, IMPRES score, as well as scores for cytotoxic cells (Figure 4a). Following that, we used CIBERSORT transcriptomic deconvolution to measure the relative abundances of 22 different immune cell types in the tumour microenvironment and compared the abundance of each cell type between the two groups. We found significant differences between the two groups across all 22 immune cell types (Figure 4b). Interestingly, immune cell types associated with the adaptive immune system such as CD4+ memory T-cells, CD8+ T-cells, and B-cells had a higher relative abundance in Group 1, whereas Group 2 had a higher relative abundance of cells typically associated with the innate immune system, such as M2 macrophages and mast cells (Figure 4b). Moreover, Group 1 samples were found to have higher abundance of lymphoid lineage cells, including B cells, CD8+ T cells and CD4+ T cells, whereas immune cells which were more abundant in Group 2 were mostly from the myeloid lineage, such as neutrophils and mast cells. There was also a notable difference in the prevalence of macrophage phenotypes between Group 1 and Group 2. Pro-inflammatory M1 macrophages were found to be more abundant in Group 1, while non-activated M0 macrophages and anti-inflammatory M2 macrophages are more abundant in Group 2 (Figure 4b).

**Figure 4.**
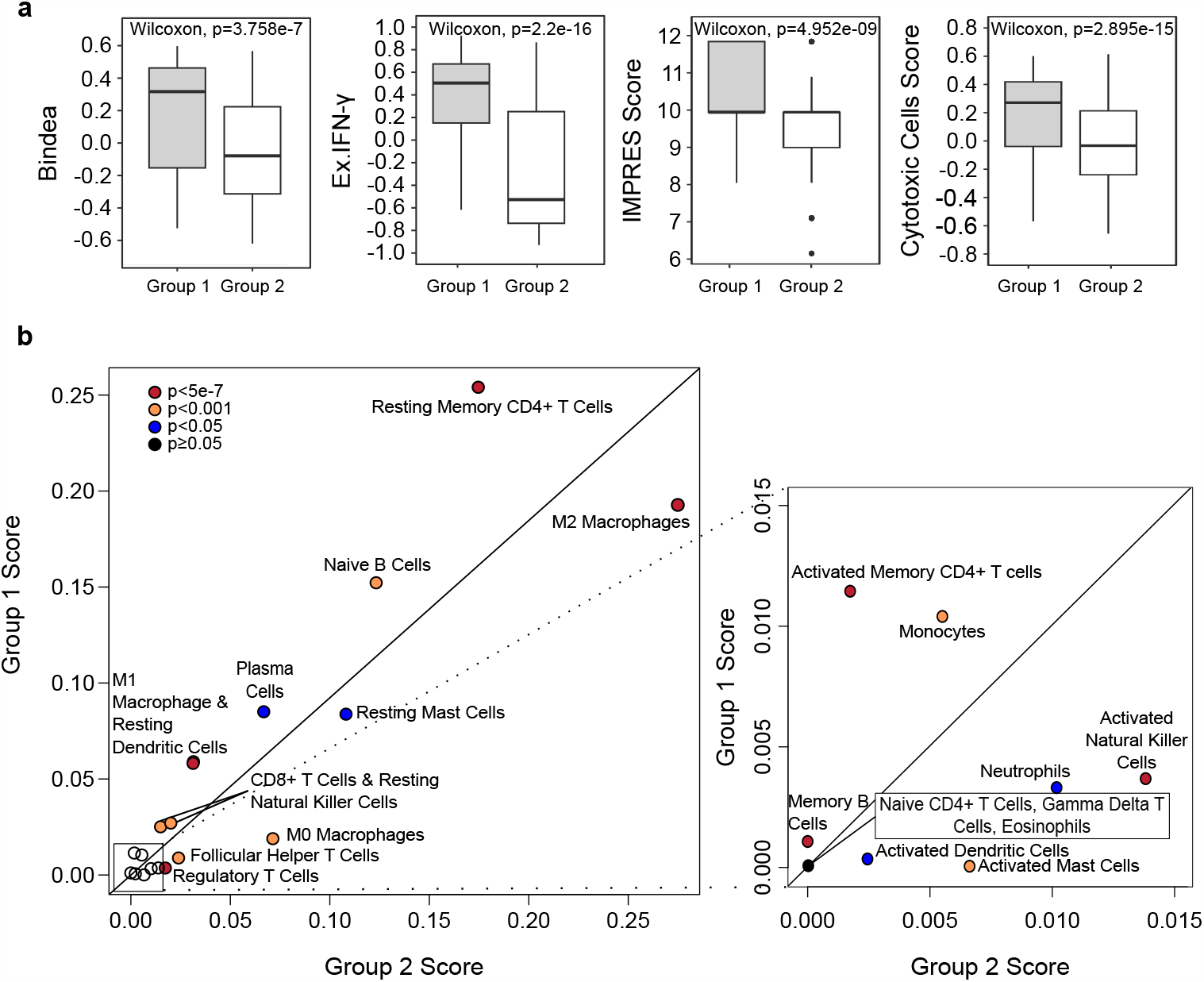
Comparison of immune profiles of Group 1 and Group 2. **(a)** Scores of Group 1 and Group 2 for BINDEA gene set, Ayers expanded IFN-gamma gene set, IMPRES score and scores for cytotoxic cells. **(b)** Comparison of abundance of immune cell types in the tumour microenvironment in Group 1 and Group 2.

### Survival Analyses of MyBrCa Clusters

Following that, we looked for prognostic differences between our clusters by conducting survival analyses. Survival analyses were performed using Cox proportional hazard models for overall survival, adjusting for tumour stage and grade. In general, HR+ clusters, (Clusters 2, 4, 5 and 7) appeared to have better survival than the rest of the cohort (Figure 5d, Supplementary Figure 4a). As expected, clusters comprising of mainly TNBC subtypes, namely Clusters 1, 8 and 11, were associated with the worst survival, followed by Cluster 3 which comprises of a mix of clinical subtypes which are mainly HER2+ or TNBCs.

**Figure 5.**
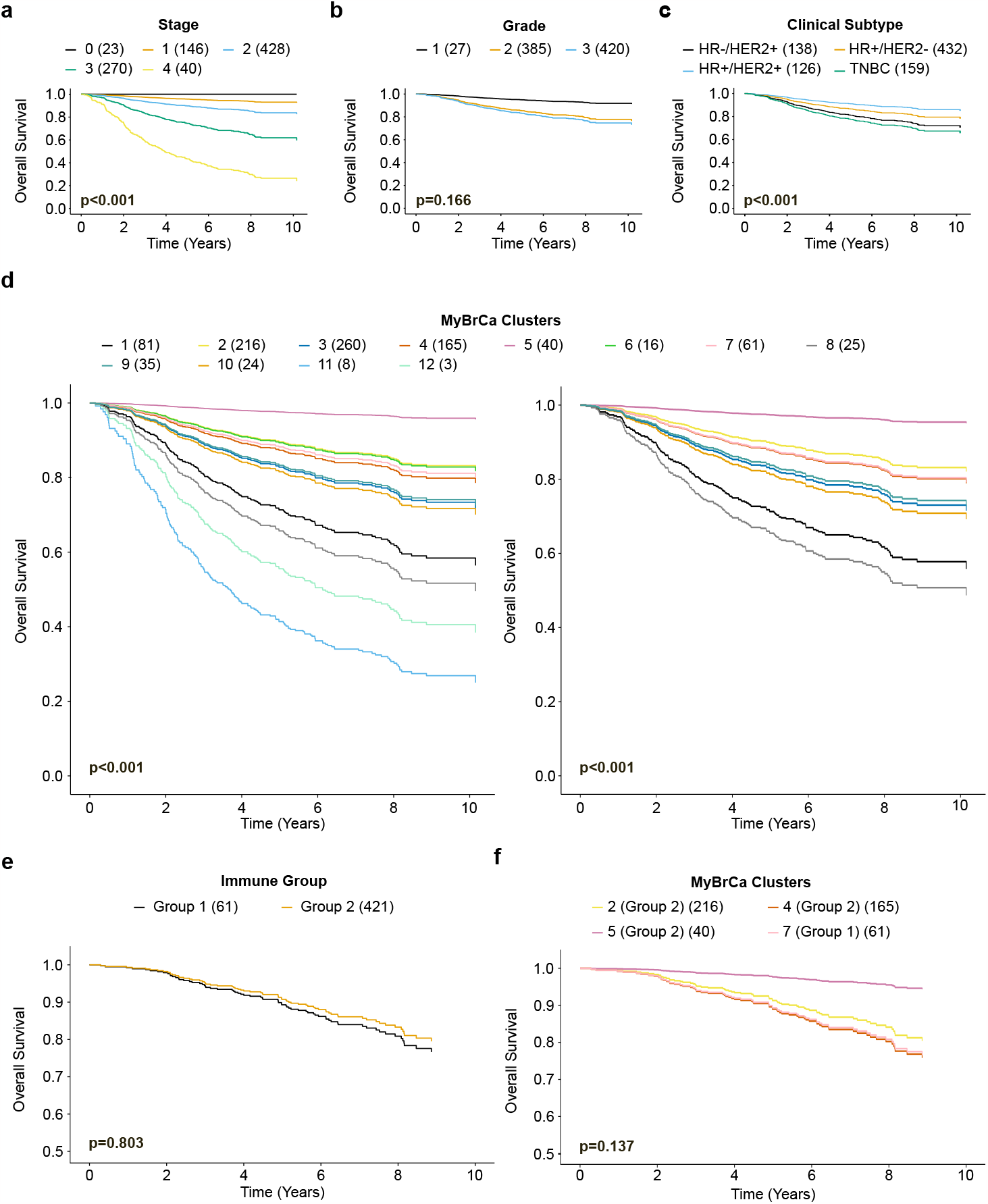
Prognostic analyses between different clusters **(a)** Overall survival by stage, adjusted for grade and clinical subtypes. **(b)** Overall survival by grade, adjusted for stage and clinical subtypes. **(c)** Overall survival by clinical subtypes, adjusted for stage and grade. **(d)** Overall survival by MyBrCa clusters, adjusted for stage and grade. In the right panel, small clusters with n<20 were removed. **(e)** Overall survival by immune group, adjusted for stage and grade. **(f)** Overall survival by MyBrCa clusters adjusted for stage and grade, showing only clusters in Group 1 or Group 2. Adjustments were made using a Cox proportional hazard model. Sample sizes are reported in brackets. For **(C)** and **(e)**, p values between the group with the best survival and the group with poorest survival are reported. For**(d)**, p values were reported for Clusters 1 and 2. For **(f)**, the p value between Cluster 7 and Cluster 5 was reported.

We also asked if survival differs between the HR+/HER2-clusters with distinct immune phenotypes (Group 1 versus Group 2). Group 1 was associated with a slightly poorer survival than Group 2, but the difference was not statistically significant (Figure 5e, Supplementary Figure 4b). Relative to Cluster 7 (Group 1), both Clusters 2 and 5 (Group 2) had better survival, although the difference was insignificant, while Cluster 4 (Group 2) had similar survival as Cluster 7 (Figure 5f, Supplementary Figure 4c).

## Discussion

The aim of this study was to look for novel subtypes of breast cancer from cluster analyses using transcriptomic data from a large Malaysian cohort of 934 breast tumours. Our classifier grouped these samples into 12 clusters which were highly associated with clinical subtypes. Comparisons of our clustering results to the PAM50 and IntClust classification schemes in the MyBrCa cohort suggest that intrinsic subtypes of breast cancer are largely conserved between Asian and Western cohorts for HER2+ and TNBC subtypes. Survival analyses of our k-means clusters were also consistent with what might be expected based on the clinical subtypes which each cluster comprise of and also consistent with previous findings. On the other hand, our results from both k-means clustering and hierarchical clustering demonstrated a unique clustering pattern of Asian HR+/HER2-breast cancer samples. Instead of conforming to the conventional luminal A-luminal B paradigm, differences in immune-related pathways, immune scores as well as immune profiles suggest that clustering of these samples were driven by immune phenotypes.

This observation suggests that in Asian populations with HR+/HER2-breast cancer, the activation of the immune system may play a more crucial role compared to other populations. These findings are consistent with previous studies which reported a more immune-active tumour microenvironment in the Asian population, supporting the notion that immune responses could potentially have a heightened importance in Asian HR+/HER2-breast cancer patients^15,36,37^. These findings are also consistent with previous studies reporting that luminal breast cancers in Asians can be further stratified based on immune profiles^38,39^. Additionally, comparison of the clustering results of our study to Integrative Clusters suggests that the majority of our HR+/HER2-immune high group (Group 1) also classified as IntClust 4, which was previously identified as having strong immune signatures in the METABRIC cohort^3^, further supporting our findings.

Previous studies have suggested that the immune activity in HR+/HER2-breast tumours is associated with different outcomes compared to other breast cancer subtypes^40^. For example, the abundance of tumour-infiltrating lymphocytes (TILs) is associated with worse outcomes in HR+/HER2-breast cancer, whereas it is associated with better outcomes in other subtypes^41^. This difference may be due to a different balance of immune cell subsets within the tumour microenvironment – such a higher abundance of FOXP3+ T-cells^42^, or due to differences in the tumour immune microenvironment such as CTLA-4 expression^43^. In addition to an association with poorer prognosis, the presence of TILs in HR+/HER2-breast cancer is also thought to be an independent predictor of response to neoadjuvant chemotherapy^44^.

The underlying factors driving the increased activation of the immune system in Asian breast cancer remains to be elucidated, however we suggest that this may be due to a combination of genetic and environmental factors. Genetic variants that lower the threshold for immune activation during breast cancer may be more common in Asian populations – for example, germline deletion of the *APOBEC3B* gene, a cytidine deaminase that has roles in both cancer mutagenesis and innate immunity, is much more common in Asian and Oceanic populations than in Western populations^45,46^. Additionally, lifestyle factors such as parity and BMI are known to influence cancer immunity^47,48^ as well as risk for developing specific breast cancer subtypes^7,8^, and these lifestyle factors also differ between Asian and Western populations, with Asian populations generally having higher (albeit decreasing) parity^49,50^, and lower BMI^51^ compared to their Western counterparts. Additionally, immunity generally declines with increasing age, and Asian breast cancer studies so far have indicated that the average age at diagnosis is lower than in Western populations^52,53^, although this may be due to a birth cohort effect^52,54^. As a whole, these studies suggest plausible mechanisms or combinations of mechanisms by which the immune system in Asian breast cancer is more active.

Our study also revealed associations between the cluster of Asian HR+/HER2-breast cancers exhibiting high immune scores and several molecular characteristics. Specifically, we observed a higher prevalence of *TP53* somatic mutations within this cluster, suggesting that reduced functionality of this well-known tumour suppressor gene may contribute to immune activation in Asian HR+/HER2-breast cancer. Interestingly, despite the strong immune activation, these Asian HR+/HER2-breast cancers demonstrated lower overall numbers of somatic SNVs and indels, as well as fewer CNAs and lower HRD scores. This is opposite to what we might expect given the current paradigm for immune activation in cancer, where high tumour mutational burden generates a high neoantigen load, leading to immune activation via tumour antigen presentation^55,56^, suggesting that immune activation in this specific subset of Asian HR+/HER2-breast cancers may be governed by non-canonical mechanisms.

In our study, the cluster of Asian HR+/HER2-breast cancers characterized by high immune scores also displayed a notable association with specific immune cell types linked to adaptive immunity. This cluster exhibited a higher prevalence of immune cell populations associated with antigenic presentation and recognition pathways, suggesting a potential mechanism of immune activation in these tumours. Furthermore, we observed a lower number of immuno-suppressive M2 macrophages within this cluster, further supporting the notion of an immunologically active tumour microenvironment. These findings are consistent with the involvement of adaptive immune responses in promoting antitumor immunity in Asian HR+/HER2-breast cancer.

In conclusion, our classifier subtyped Asian breast cancer into biologically distinct clusters which revealed that the clustering of Asian HR+/HER2-breast cancer is driven by immune phenotypes. These findings may be important because HR+ breast cancer is traditionally associated with a less immune active tumour microenvironment and thus less responsive to immunotherapy, but the findings from this study support other recent studies suggesting that a subset of HR+/HER2-breast cancer may be more likely to respond to immunotherapy.

## Abbreviations list

MyBrCa: the Malaysian Breast Cancer cohort
HRD: homologous recombination deficiency
TNBC: triple negative breast cancer
RNA-seq: RNA sequencing
CNA: copy number aberration
GSVA: gene set variation analysis
SBS: Single Base Substitutions
SNV: single nucleotide variation
GSEA: gene set enrichment analysis
IFN-γ: interferon gamma

## Acknowledgements

Cancer Research Malaysia receives charitable funding from the Scientex Foundation, Estée Lauder Companies, Vistage Malaysia, Yayasan PETRONAS, and Yayasan Sime Darby which contributed to the funding of this study. Funding was also provided by a research grant from the Newton-Ungku Omar Fund (MRC Ref: MR/P012442/1) to SFC and SHT. OMR, CC, and SFC also receive funding from Cancer Research UK. All genomics work was undertaken by the Genomics Core Facility CRUK Cambridge Institute.

## Author Contributions

JWP, MR and SHT contributed to the experimental design, data analysis, supervised experiments, wrote the manuscript and generation of figures. WQC contributed to data analysis, writing of manuscript and generation of figures. SNH, TI, LYT, SJ, MHS, CHY, PR, LML, and AMT contributed to sample collection and processing and data collection, while OMR and SFC generated and processed sequencing data. PR and LML provided histopathology expertise, and collected clinical data together with MHS, TI, SJ, LYT, CHY and AMT. CC, SFC and SHT contributed to obtaining funding for the project. The work reported in the paper has been performed by the authors, unless clearly specified in the text.

## Data Accessibility

The whole exome sequencing and RNA-seq data included in this study are available in the European Genome-phenome Archive under accession numbers EGAS00001006518 and EGAS00001004518. Access to controlled patient data will require the approval of the Data Access Committee. Further information is available from the corresponding author upon request.

## Ethics Declaration

Patient recruitment and sample collection was reviewed and approved by the Independent Ethics Committee, Ramsay Sime Darby Health Care (reference no: 201109.4 and 201208.1), as well as the Medical Ethics Committee of the University Malaya Medical Centre (reference no: 842.9). Written informed consent to participation in research was given by each individual patient.

