## Supplementary Figure 1 to 4 for "Clustering of HR+/HER2- breast cancer in an Asian cohort is driven by immune phenotypes"

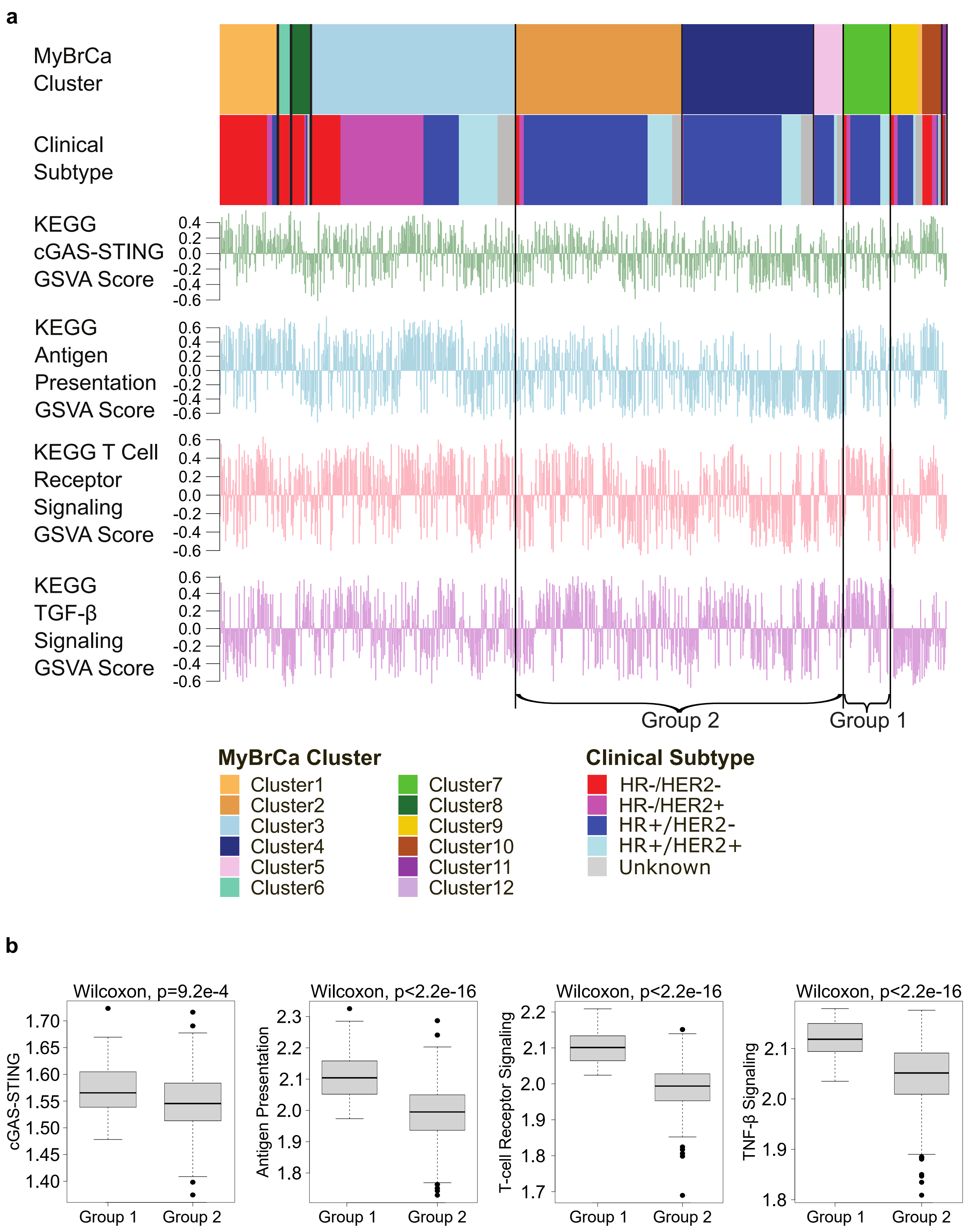


**Supplementary Figure 1 (a)** Comparison of MyBrCa Clusters and clinical subtypes with KEGG pathway GSVA scores. HR+ clusters with high immune scores (Group 1) and low immune scores (Group 2) are indicated. **(b)** Comparison of KEGG pathways GSEA scores between Group 1 and Group 2.

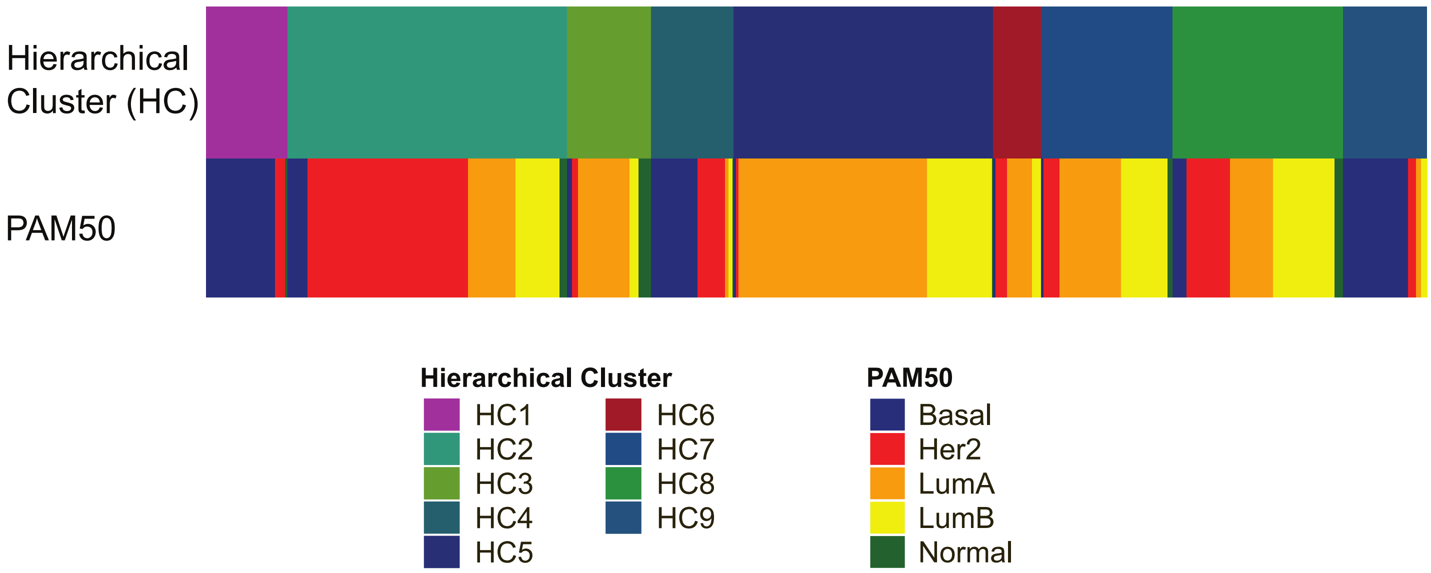


**Supplementary Figure 2** Comparison of clustering results of the MyBrCa cohort using hierarchical clustering (HC) and PAM50.

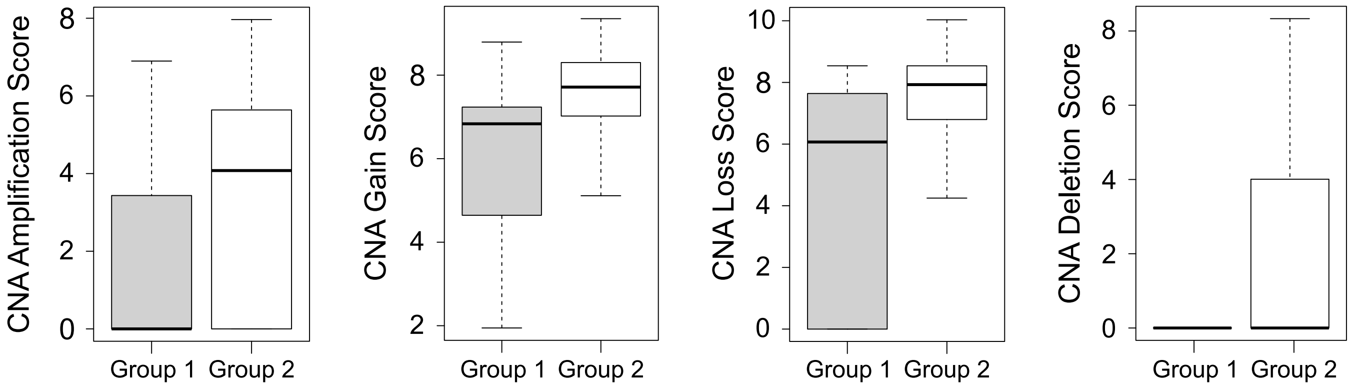


**Supplementary Figure 3** Comparison of Copy Number Aberration (CNA) amplification, gain, loss and deletion scores between Group 1 and Group 2.

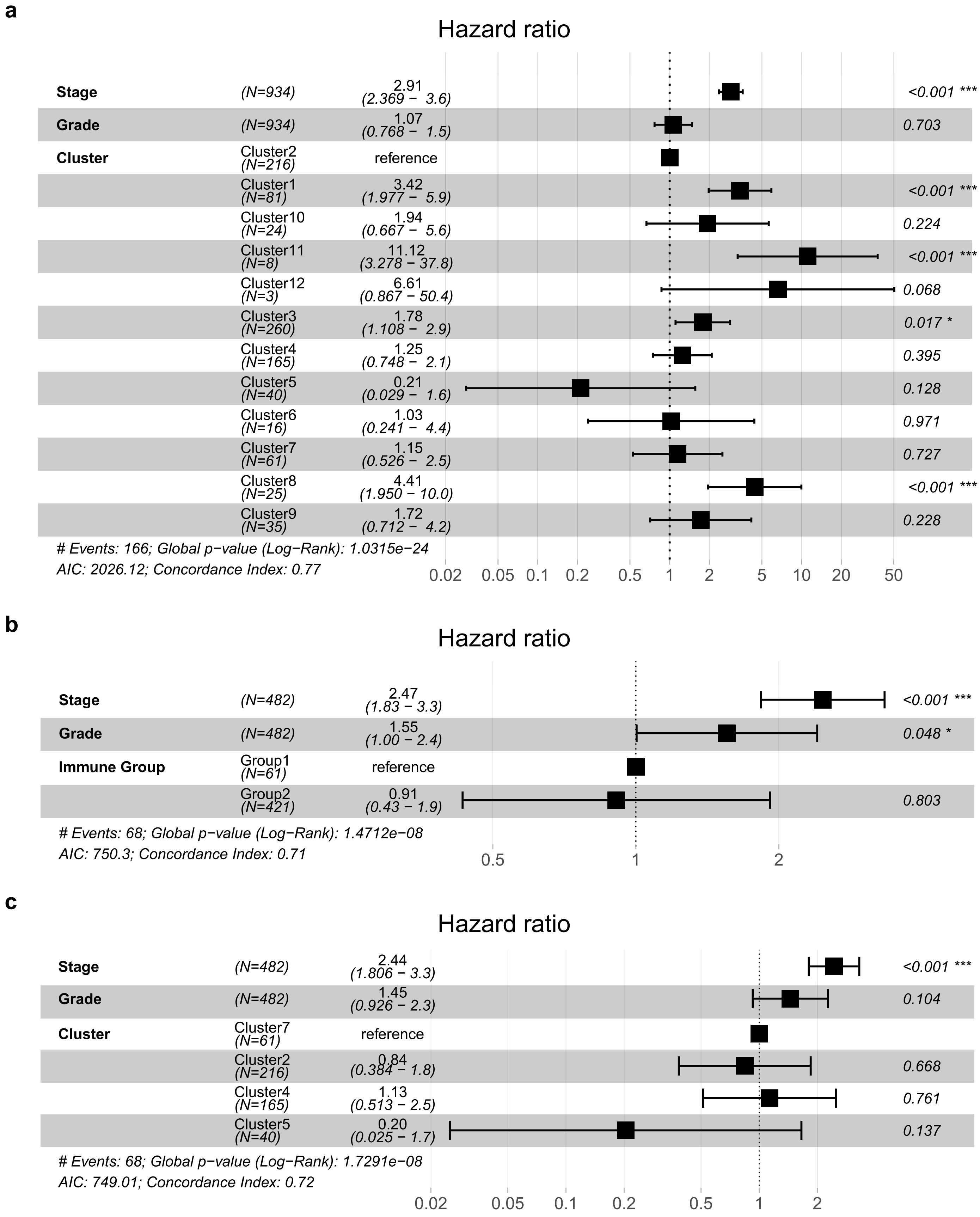


**Supplementary Figure 4 Cox proportional hazard models of overall survival of MyBrCa patients. (a)** Stage, grade and clusters were included as variables. **(b)** Stage, grade and immune group were included as variables. **(c)** Stage, grade and clusters were included as variables, but only clusters belonging to Group 1 and Group 2 were included. Error bars represent 95% confidence interval of hazard ratio.
